# Distance-AF: Modifying Predicted Protein Structure Models by Alphafold2 with User-Specified Distance Constraints

**DOI:** 10.1101/2023.12.01.569498

**Authors:** Yuanyuan Zhang, Zicong Zhang, Yuki Kagaya, Genki Terashi, Bowen Zhao, Yi Xiong, Daisuke Kihara

## Abstract

The three-dimensional structure of a protein plays a fundamental role in determining its function and has an essential impact on understanding biological processes. Despite significant progress in protein structure prediction, such as AlphaFold2, challenges remain on those hard targets that Alphafold2 does not often perform well due to the complex folding of protein and a large number of possible conformations. Here we present a modified version of the AlphaFold2, called Distance-AF, which aims to improve the performance of AlphaFold2 by including distance constraints as input information. Distance-AF uses AlphaFold2’s predicted structure as a starting point and incorporates distance constraints between amino acids to adjust folding of the protein structure until it meets the constraints. Distance-AF can correct the domain orientation on challenging targets, leading to more accurate structures with a lower root mean square deviation (RMSD). The ability of Distance-AF is also useful in fitting protein structures into cryo-electron microscopy maps.

## Introduction

Protein structure prediction is an important and challenging problems. Prior to Critical Assessment of Protein Structure Prediction (CASP13), traditional methods like I-Tasser [1], FragFold [2] are based on fragments derived from homologous structures to predict protein structures. With the development of deep learning, methods such as CONFOLD2[3], RaptorX[4] have emerged, which improved the prediction performance. Furthermore, notable methods like AlphaFold [5], developed for CASP13, have shown impressive performance. This progress was further amplified in CASP14 [6], where AlphaFold2[7] and RoseTTAFold[8] made further improvement in the modeling accuracy. AlphaFold2 is a deep learning-based model that has gathered significant attention due to its notable performance in CASP14. It outperformed other methods and achieved unprecedented accuracy in predicting protein structures. Its ability to accurately capture the intricate folding patterns of proteins has been exemplified for better understanding protein function and aiding drug discovery efforts.

Despite its impressive performance, it is important to note that AlphaFold2 is not always successful. It still encounters challenges in predicting the structures of certain proteins. The complex nature of protein folding, with its intricate energy landscapes and conformational diversity, presents inherent difficulties even for AlphaFold2. AlphaFold2 performs proficiency in accurately predicting individual domain structure, it may encounter challenges in accurately determining domain orientations. Therefore, further improvements are required to enhance its ability to correctly predict the spatial arrangement and orientations of these domains within the overall protein structure.

Here, we propose Distance-AF, a novel deep learning-based approach that builds upon the structure module of AlphaFold2 by incorporating additional information of distances between amino acids. By considering the precise distances between amino acids, we exploit the inherent spatial relationships present in proteins and construct a comprehensive distance matrix. This matrix plays a vital role in accurately predicting the intricate 3D structures of proteins within the Distance-AF framework. By utilizing limited number of distance constraints and taking the single and pair embeddings from AlphaFold2, Distance-AF effectively captures the geometric relationships between amino acids and significantly enhances the accuracy of predicted protein structures obtained solely from AlphaFold2 in an iteratively optimization way. Through rigorous evaluation and in-depth analysis, we show the remarkable performance of Distance-AF compared to existing methods on multiple evaluation scores on 25 targets. Furthermore, to demonstrate the wide practical applications of Distance-AF, our evaluation encompasses diverse protein targets, including the challenging category of G-protein coupled receptors (GPCRs), targets derived from Cryo-EM maps and multiple ensembled NMR targets.

## Methods

### Overview of Distance-AF

AlphaFold2 leverages search mechanism against clustered protein sequences databases to gather information from multiple sequences alignment. The sequential information is encoded into both single and pair representations using the Evoformer module. The single representation contains encoding at the residue level, while the pair representation embeds geometric distance information between pairs of amino acids. Built upon AlphaFold2, for a given sequence, Distance-AF takes single and pair representations, distance constraints as input, and predicts the 3D protein structure. In general, it operates in 2 steps. First, single and pair representations pass through structure module, to complete 3D structure prediction, which is regarded as the initial structure; second, distance constraints are incorporated to enhance the accuracy of the 3D structure prediction on the top of initial structure. This is achieved by calculating the loss function, which quantifies the discrepancy of the computed distances between amino acids in the predicted protein structure and the provided distance constraints. This information is iteratively backpropagated to optimize the parameters in the neural network in of IPA module, ensuring that the predicted structure adheres to the given distance constraints. The framework of Distance-AF is illustrated in Fig 1.

**Fig 1.**
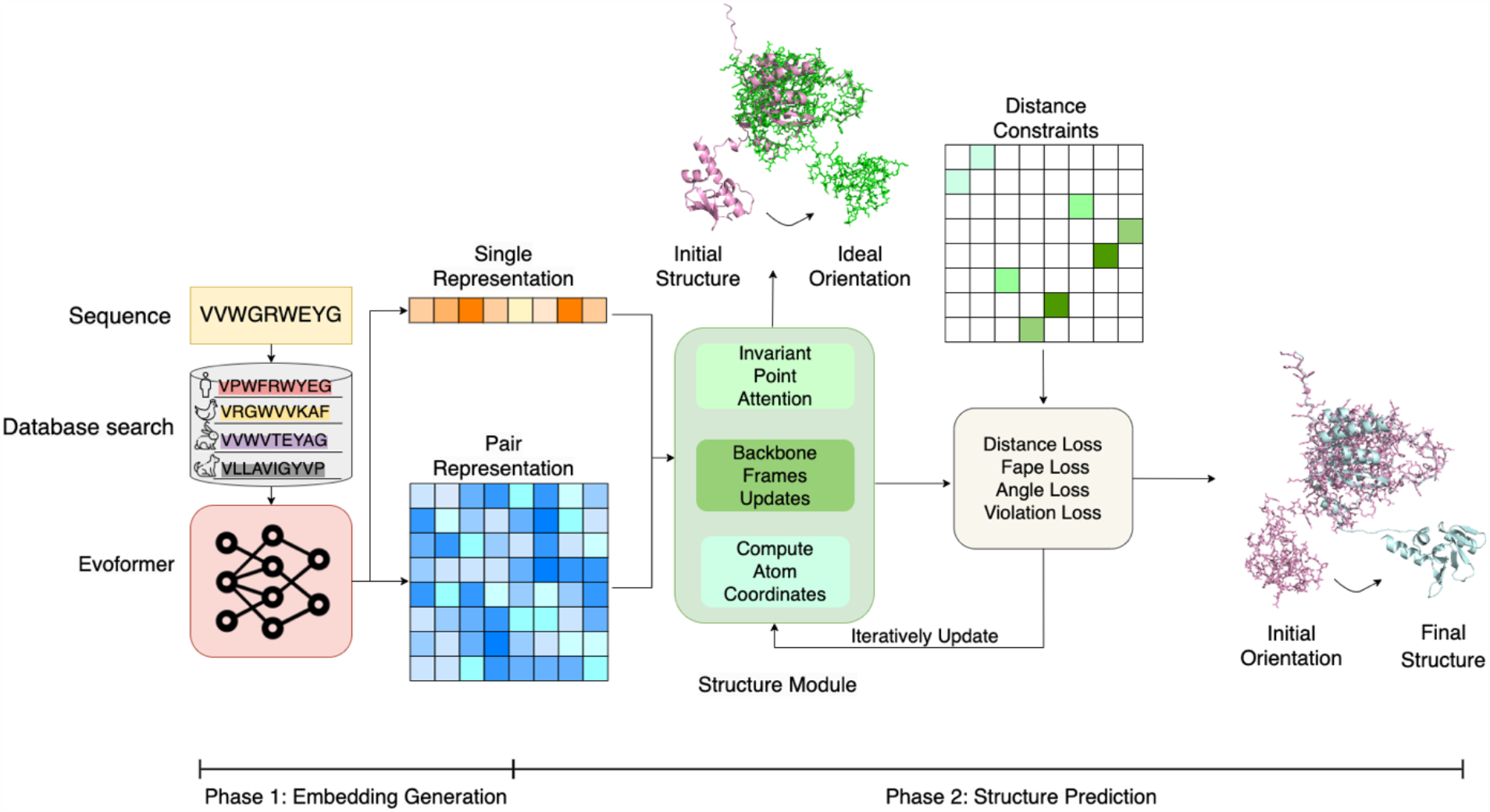
Overall framework of Distance-AF consists of two phases. In Phase one, the method takes the protein sequence as input and performs multiple sequence alignment (MSA)[9] and template search against databases. The MSA and template features are then embedded and paired into two embeddings, MSA and pair representations. The evoformer with axial self-attention layers, utilizing frozen weights from AlphaFold2, processes the representations and generates single representation and pair representation as the final output of Phase one. Phase two of Distance-AF begins with inputting the two representations obtained from Phase one into the structure module. This module updates the backbone frames and single representation using invariant point attention layers and predicts atom-level 3D coordinates. To refine the predicted structure, distance constraints are introduced, which are defined as Euclidean distance on pairs of residues at the Cα atoms. Subsequently, the backbone is adjusted to meet these distance constraints. Once the moved backbone satisfies the constraints, the restoration of the sidechain structure is initiated using the local initial structure and ultimately leading to improved predictions that align closely with the native structure.

### Representations generation

To obtain single and pair representations, we utilize the AlphaFold2 framework. The general process in AlphaFold2 begins by inputting a protein sequence, which undergoes a database search. This search provides valuable insights into the co-evolutionary patterns of amino acids. The subsequent evoformer layers, equipped with an attention mechanism, learn and encode residue interactions into representations. Finally, the framework outputs the single and pair representations. One notable advantage of our pipeline is the consideration of the substantial time required to generate MSAs. We employ MSA search on the Uniref30 database[10] instead of searching the full database (Mgnify[11], Uniref90[12], Uniclust30[10], and BFD) within AlphaFold2, which saves much time.

### Distance constraints definition

For specific protein structures, the desired distance constraints used by Distance-AF are defined as the expected Euclidean distances between pairs of *Cα* atoms at amino acids. These distance constraints can be derived from experimental methods like cryo-electron microscopy (cryo-EM)[13], nuclear magnetic resonance spectroscopy (NMR)[14], or chemical cross-linking[15]. In Distance-AF, distance constraints serve as important guidance during the modelling process, improving the accuracy and reliability of the predicted 3D structure. They ensure that the generated structures adhere to the experimentally observed spatial relationships between amino acids.

### Iteratively overfitting on Distance-AF

Distance-AF employs an overfitting mechanism, iteratively updating network parameters until the predicted structure satisfies the given distance constraints. This iterative process allows the model to fine-tune its predictions, aligning them more accurately with the actual distances between amino acids. Thus, for various targets, Distance-AF eventually generates diverse sets of parameters tailored to specific targets, enabling it to effectively capture the unique characteristics of each protein. The whole procedures include the following steps, first, single and pair representation are as input into structure module, which predict the 3D protein structure; second, the coordinates of the *Cα a*toms corresponding to given distance constraints are predicted; next, the predicted distances between pairs of *Cα* atoms are regarded as the divergence between the predicted and native structures by comparing to given distance constraints. To minimize the divergence and achieve the goal that predicted distances are supposed to be as close as possible to native distances, we define a new loss function called distance loss as Eq 1. shows:

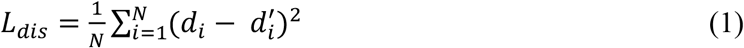

Where *d*_*i*_ is the native distance on the pair of *Cα* atoms, 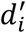 is measured distance on predicted structure at the same pair of *Cα* atoms. *N* is the number of distance constraints. *L*_*dis*_ is integrated into final loss function with various weights during different optimization stages.

The final loss function comprises four components. The first part is the distance loss in Eq 1., which involves iteratively adjusting the positions of *Cα* atoms to satisfy the specified distance constraints. The second component is the modified Frame aligned point error (FAPE) loss from AlphaFold2[7], which prevents the local structure from being destroyed during the optimization process driven by the distance loss function. This is necessary as the local 3D structure may be compromised when the network is optimized solely for distance constraints. The third component is the angle loss, referring to the loss function that penalizes the deviation of predicted torsion angles, including *φ, ψ, ω* for backbone and *X*_*1*_, *X*_*2*_, *X*_*3*_, *X*_*4*_ dihedral angles at sidechain. The fourth is the violation loss, related to structural violation in the predicted structure. Violation loss penalizes unidealized bond lengths, peptide bond angles and atom clashes, the distance between non-boned atoms inconsistent with physical properties[7]. In Distance-AF, as the true positions are unknown, we take the original structure predicted by AlphaFold2 as the pseudo-true positions in FAPE, angle and violation loss. Considering the various consequences of whether distance constraints are satisfied for the overall network optimization, Distance-AF uses dynamic weights to reweight each loss function at various phases. Eq. 2 shows the details about loss function.

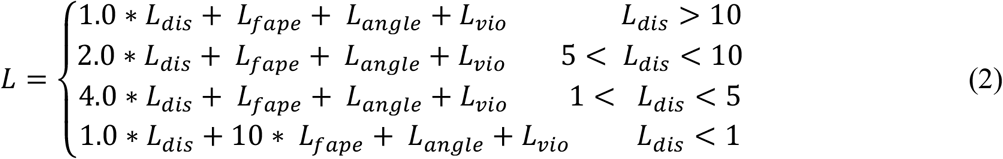

In Eq.2, the weights of *L*_*dis*_ *a*nd *L*_*fape*_ *c*hange flexibly with the value of *L*_*dis*_. When *L*_*dis*_ *i*s greater than 10, which taking a large portion dominates the backpropagation process to optimize it. Thus, we assign it a small weight to avoid unbalanced parameter updates leading to local structure distortion. When *L*_*dis*_ is at the range of 1 to 10, the distances in prediction are still unsatisfied to given constraints, so we add some weights to *L*_*dis*_, expecting the neural network to still focus on optimizing distance loss. While *L*_*dis*_ achieves to ideal values, less than 1.0, meaning the predicted distances are almost satisfy given distance constraints, we lower down the weights of *L*_*dis*_ and emphasize on *L*_*fape*_, with the goal that more efforts in backpropagation lead to local structure reconstruction in avoidance of too strict distance consistency at current stage.

In particular, when utilizing Distance-AF for structure prediction on targets with defined distance constraints spanning distinct domains, it becomes imperative to address potential domain motion. In these instances, the FAPE (Frame Aligned Point Error) loss within the Distance-AF framework is adapted to exclusively calculate loss for residues within the same domain.

Domain motion encompasses the structural shifts or relative displacements between discrete domains within a protein which holds substantial impact over the overall protein structure and its functional kinetics. However, when combined with the distance loss in Eq1, FAPE loss introduces a counterproductive effect by incorporating the loss between residues from different domains, effectively constraining the movement and compelling it back to its initial position. To resolve this problem, the formulation of FAPE loss within the Distance-AF framework is customized to specifically compute loss for interactions occurring within the confines of a single domain. This adjustment in the training process places greater emphasis on accurately situating and assembling fragments within each domain independently, all the while preserving the capacity for domain motion and flexibility.

Overall, the modification of the FAPE loss in the Distance-AF framework to compute loss for intra-domain interactions when distance constraints are defined on different domains represents a crucial adaptation. To achieve the modified version of FAPE loss, the domain separation will be passed to Distance-AF, this allows Distance-AF to differentiate between different domains and treat them as separate components by masking the parts related to inter-domain loss computation. The detailed pseudo-code of modified FAPE loss is as Fig 2. shows.

**Fig 2.**
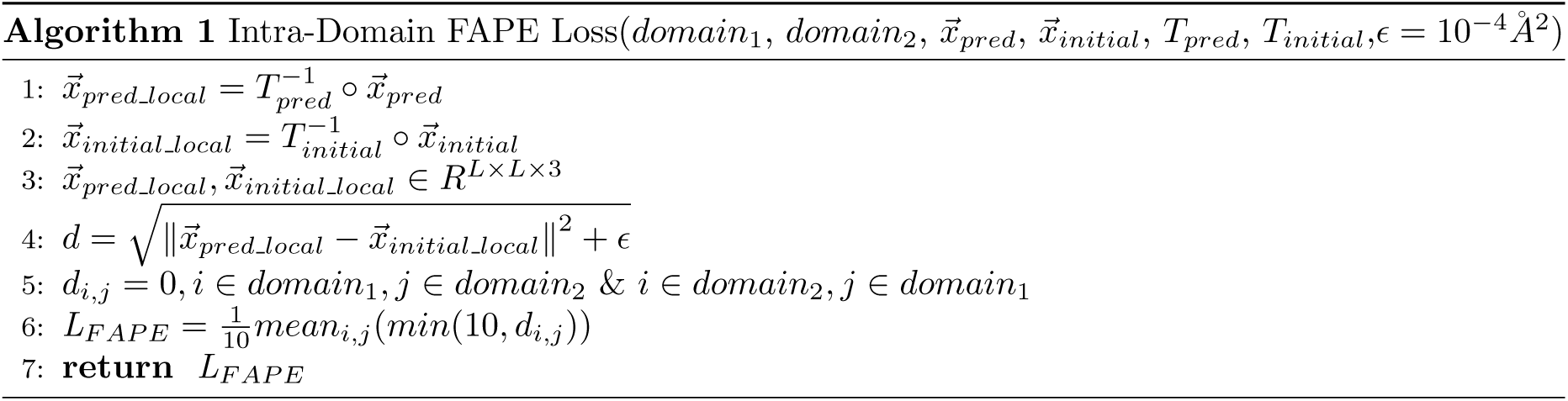
The algorithm of modified FAPE loss only considering the residues belonging to intra-domain. As it is shown in line 3, those values calculated but associated with amino acids from different domains would be masked as 0 to avoid pulling the domain movement back achieved by distance loss.

For other network settings of training Distance-AF, the network depth of structure module is 8, the learning rate is 0.001 using the Adam optimizer with *β*_*1*_ *= 0*.9 and *β*_*2*_ *= 0*.99, controlling the exponential decay rate for the moving average of the first-order moment (mean) and second-order moment (variance) of the gradients.

### Evaluation metric

To evaluate the performance of Distance-AF, several evaluation scores are considered. These include Full-atom RMSD (Root Mean Square Deviation)[16], which measures the similarity between the predicted protein structure and the native structure at the level of individual atoms. The TM-score[17], which ranges from 0 to 1, is based on structural template comparison and calculates the RMSD between the aligned Cα atoms of the two structures. It is then normalized by a scaling factor that accounts for the length of the protein. Another criterion is GDT-TS[18], which assesses similarity by measuring the percentage of residues in the predicted structure that fall within specific distance thresholds compared to the native structure. These distance thresholds are typically set at 1, 2, 4, and 8 Å. A higher GDT-TS score indicates that the predicted residues are closer to the reference residues. Additionally, GDT-HA[18] calculates the percentage of residues in the predicted structure that fall within stricter distance thresholds compared to the reference structure. These distance thresholds are typically set at 0.5, 1, 2, and 4 Å. The aim of GDT-HA is to identify residues in the predicted structure that are extremely close to the reference structure, reflecting a high degree of accuracy.

## Results

To demonstrate the performance of Distance-AF, we initially benchmarked it against 25 targets. This benchmarking process involved comparing Distance-AF to AlphaFold2[7] and Rosetta[19] using the four criteria above. Following the benchmarking, we provide the visualization of examples predicted by Distance-AF, which serve to highlight the model’s accuracy and high-quality predictions. Finally, we explore the implications of Distance-AF within the realm of GPCRs (G-protein coupled receptors)[20], Cryo-EM deposited and NMR targets with ensembles[21].

### Dataset construction

Thirty-eight protein monomers are selected from the Protein Data Bank (PDB)[22] for benchmarking and application purposes. Among these, twenty-five are normal monomer entries as general evaluation, further for application assessment, five are GPCRs, five are associated with EMDB, and the rest 3 are protein ensembled targets. The selection of targets follows specific criteria. For normal monomer entries, first, all targets are searched from the AlphaFold2 Database[23], where AlphaFold2 is unable to predict accurately with RMSD values greater than 10 Å, second, these targets are filtered based on their average pLDDT score in descending order, ensuring a high quality of local structure prediction by AlphaFold2. Thirdly, most of the selected targets contain two distinct domain regions, aiming to correct any shifted domain causing a larger RMSD using Distance-AF. For GPCR cases, as the conformation change differs slightly between active and inactive states with much smaller RMSD, we select those whose structural rearrangement is folded incorrectly visually. For targets associated with Cryo-EM maps, those with predicted structures by AlphaFold2 falling outside the volume are considered. The last three targets are structural ensembles of intrinsically disordered proteins (IDPs) chosen from Protein Ensemble Database (PED)[21], where the structural variations are all about domain conformations change.

### Comparison to AlphaFold2

In Distance-AF, several distance constraints are selected in advance before running. For typical targets, we choose 6 distance constraints for them. For the 25 targets under benchmarking, all of them contain 2 separate domains. So, while selecting distance constraints for them, each distance constraint connects 2 *Cα a*toms from different domains. About the strategy to choosing distance constraints, we follow 2 rules. First, considering the targets have different oriented domains, some desired distances deviate significantly from the ones in the initial predicted structure. Therefore, the selected distance constraints are expected to provide such divergent information, with the difference between distances and initial distances being as large as possible. Second, since Distance-AF aims to move the whole domains to ideal conformation in line with the given distance constraints, it is better that the geometric distribution of selected distance constraints covers a large portion of the moved domain. Following the idea of constraint selection, we sampled 6 distance constraints for the 25 targets and run with Distance-AF. The benchmarking results are shown in Fig 3.

**Fig 3.**
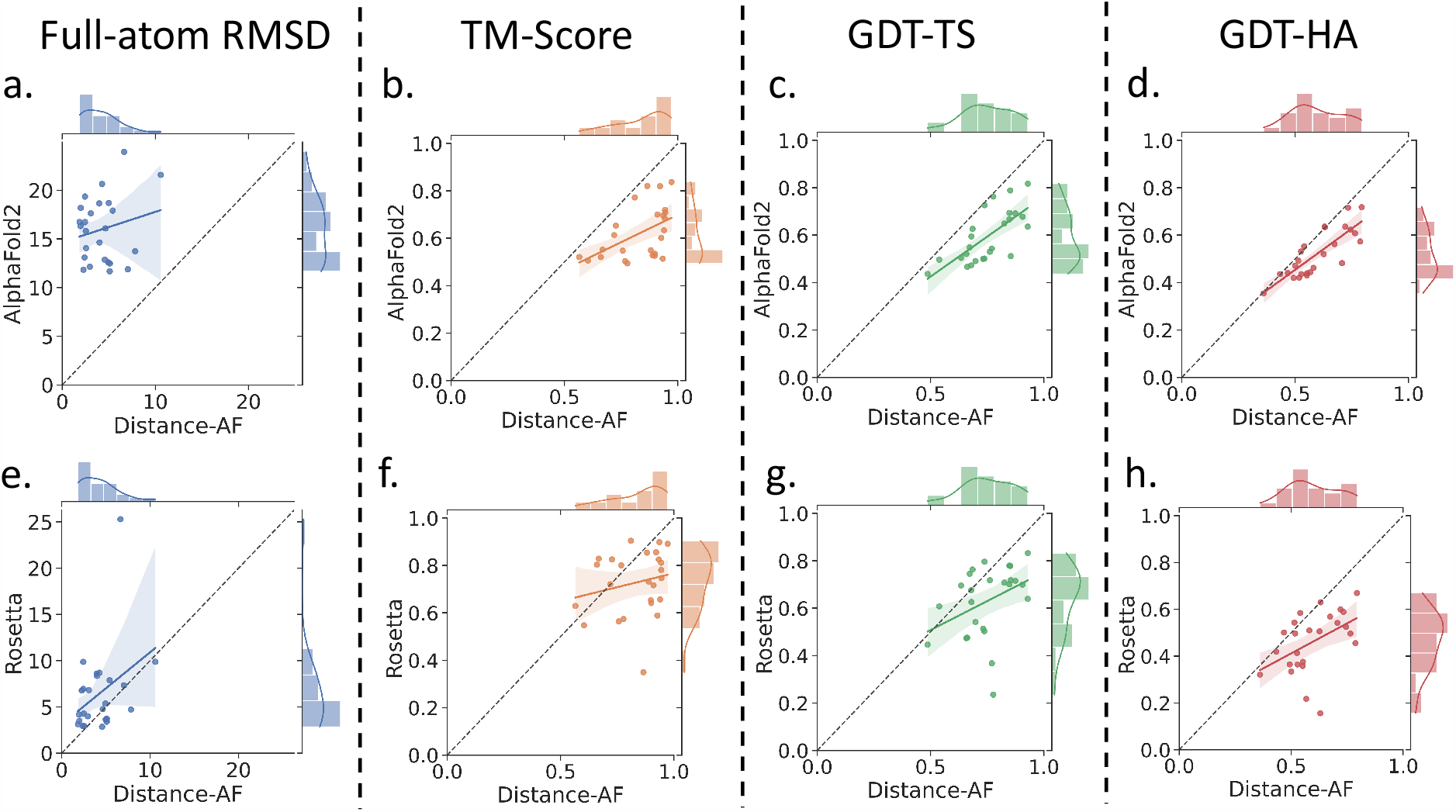
Benchmarking on the 25 monomer targets, a. comparison of Distance-AF(x-axis) and AlphaFold2 (y-axis) about RMSD; b. comparison of Distance-AF(x-axis) and AlphaFold2 (y-axis) about TM-score,; c. comparison of Distance-AF(x-axis) and AlphaFold2 (y-axis) about GDT-TS; d. comparison of Distance-AF(x-axis) and AlphaFold2 (y-axis) about GDT-HA. To summarize for a,b,c,d, Distance-AF shows superior performance on all targets than AlphaFold2 on all 25 targets at RMSD, TM-Score and GDT-TS, while AlphaFold2 slightly outperforms Distance-AF at 5 targets on GDT-HA. Next, e. comparison between Distance-AF(x-axis) and Rosetta(y-axis), Distance-AF performed better at 19 out of 25 targets than Rosetta on RMSD; f. comparison about TM-score of Distance-AF(x-axis) and Rosetta(y-axis), which shows the similar trend to RMSD, Distance-AF wins on 19 targets; g. the comparison about GDT-TS between Distance-AF(x-axis) and Rosetta(y-axis), for this metric, Distance-AF works better at 20 out of 25 targets; h. the comparison about GDT-HA between Distance-AF(x-axis) and Rosetta(y-axis), Distance-AF beats Rosetta on 22 out of 25 targets.

Fig 3 a,b,c,d depict four summarized scatter plots that are directly related to RMSD, TM-score, GDT-TS, and GDT-HA evaluations across the 25 examples on Distance-AF and AlphaFold2. These scatter plots provide a visual distribution of these metrics, by examining the patterns and trends displayed in these plots. It is evident that Distance-AF outperforms AlphaFold2 across all four evaluation scores for the 25 targets considered. To be specific, Distance-AF successfully improves the RMSDs for all 25 targets to 10 Å below, even 18 out of 25 are under 5

Å, meaning the high quality of predicted structures by Distance-AF, while AlphaFold2 produces the 25 structures with RMSD greater than 10 Å generally. As for TM-score, Distance-AF also outperforms AlphaFold2, with 16 out of 25 targets achieve TM-score more than 0.8, while AlphaFold2 has only 3 targets above 0.8. Additionally, Distance-AF also wins at GDT-TS on all 25 targets, and 15 out of 25 cases with the structures predicted by Distance-AF obtain relatively higher GDT-TS at above 0.7, compared to 4 out of 25 on AlphaFold2. For GDT-HA, Distance-AF outperforms on 20 among 25 targets compared to AlphaFold2.

### Comparison to Rosetta

Rosetta[19, 24, 25] is a powerful computational tool extensively developed for protein structure refinement, particularly in the presence of distance constraints. Its incorporation of distance constraints enhances the accuracy of protein structure predictions, guiding the conformational sampling process and allowing Rosetta to explore the vast conformational space more effectively. Leveraging energy-based scoring functions and sophisticated sampling strategies, Rosetta optimizes the agreement between predicted protein structures and experimental distance constraints, ultimately improving the accuracy and reliability of protein structures. To demonstrate the performance of Distance-AF, we run Rosetta on the 25 cases we have experimented on with the same distance constraints. The version we used is Rosetta 3.13, which is downloaded from the official website of RosettaCommons[24, 25].We considered multiple factors that may be related to Rosetta’s performance while in evaluation. First, considering that the refined structures from Rosetta diverse a lot under a variety of Monte Carlo searching directions, we allow Rosetta to generate 10 structures for each target and choose the best one with lowest energy score to compare against Distance-AF. Second, a parameter called distance constraint weight to be set manually in Rosetta also plays a vital role in final structure. To find out the most appropriate weight to produce the best structure for Rosetta, we tested 5 values for it, 0.01, 0.1, 1.0(default set by Rosetta), 5.0 and 10.0 on several targets. By checking and comparing the quality structures, eventually we decided to use 0.1 which performed best than other values generally. The performance comparison on 4 metrics has been shown in Fig 3. e, f, g, h, which specifically corresponds to full-atom RMSD, TM-score, GDT-TS and GDT-HA. For both RMSD and TM-score, there are 19 targets which Distance-AF works better than Rosetta. If we focus on those structures which are predicted with high quality, which are with RMSD less than 5 Å, Distance-AF succeeds on 18 targets against Rosetta on 13. Meanwhile, with TM-score of 0.8 as a cut-off, Distance-AF have 16 targets at this range while Rosetta only manages on 11 targets. Then for GDT-TS Distance-AF predicts more accurate structures than Rosetta on 20 out of 25 cases in terms of GDT-TS, with 15 targets get 0.7 or higher GDT-TS score, 4 more than Rosetta. Finally at GDT-HA, Rosetta works worse than Distance-AF that Distance-AF wins 22 targets. There are 7 targets with GDT-HA above 0.7 for Distance-AF, but none for Rosetta.

### Case study of predicted structures

Fig 4 shows 4 example cases. The reference structures are in green, and the predicted models are shown with different colors, AlphaFold2 is in cyan, and Distance-AF is in magenta. Chain B of 6VW7(Figure 4 a.), chain B of 1NT2 (Figure 4 b.), chain A of 1IXC (Figure 4 c.) and chain A of 2BJ7 (Figure 4 d.) are predicted almost perfectly by Distance-AF with an RMSD of 1.835 Å, 2.389 Å, 1.958 Å and 2.278 Å, respectively. AlphaFold2 struggled with the prediction but on very large RMSD values at 18.690 Å, 13.097 Å, 16.328 Å and 11.829 Å, with clear wrong domain orientation. With the given 6 distance constraints defining on residues from different domains, Distance-AF correct the wrong domain orientation into a near-native way.

**Fig 4.**
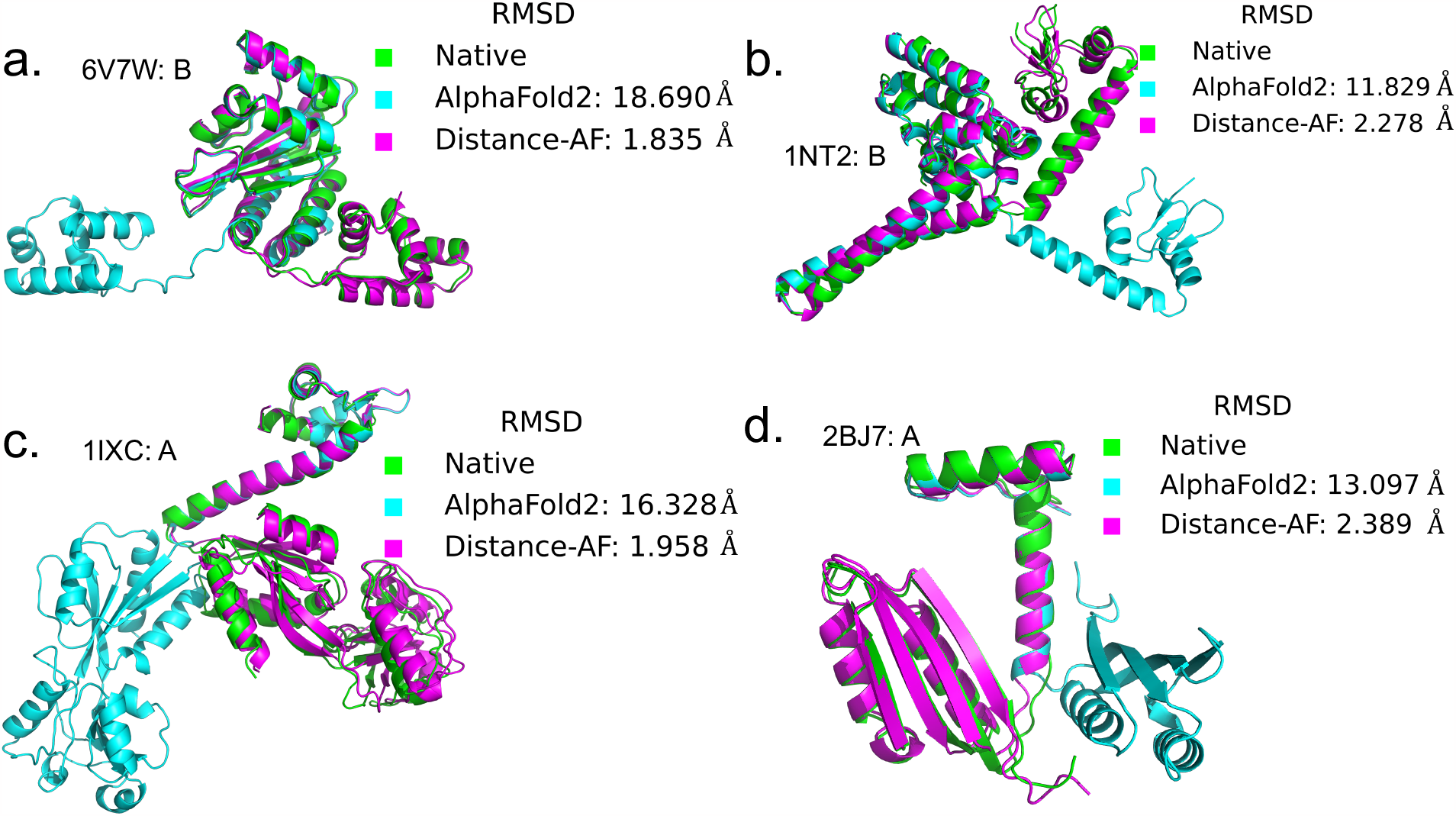
Examples for case study, whose RMSD are greater than 10 Å on AlphaFold2’s predicted structures. Distance-AF shows substantial improvement overAlphaFold2 with significantly lower RMSD and are with much higher agreement with reference structures. a. Distance-AF performs much better than AlphaFold2 by decreasing the full-atom RMSD into 1.835 Å, with a difference of 16.865 Å on the chain B of 6V7W. b. The structure predicted by Distance-AF on chain B of 1NT2, has 9.551 Å of full-atom RMSD lower than the one predicted by AlphaFold2. c. For chain A of 1IXC, Distance-AF outperform AlphaFold2 by producing near native structure with only 1.958 Å against that of 16.328 Å by AlphaFold2. d. This is an example of chain A of 2BJ7, the predicted structure by Distance-AF is more accurate by reducing the RMSD of 10.608 Å from the one by AlphaFold2.

### Implications of Distance-AF to structures deposited from Cryo-EM

We found that although AlphaFold2 generally performs well in predicting protein structures, there are instances where it produces incorrect predictions. This becomes evident when comparing the predicted structures to experimentally determined structures obtained from Cryo-EM maps. These high-resolution experimental structures serve as reliable references for assessing the accuracy of protein predictions. In order to evaluate the ability of Distance-AF to address the limitations of AlphaFold2 on these targets, we choose several such cases and run Distance-AF and then evaluate the performance.

In Figure 5, native structures are depicted in green with a cartoon representation. The structures in cyan represent predictions generated by AlphaFold2, exhibiting higher RMSD values and domain inconsistency. Conversely, structures in magenta correspond to predictions made by Distance-AF. These demonstrate improved alignment with the native structure, resulting in an enhanced overall match between the predicted structure and the cryo-EM density.

**Fig 5.**
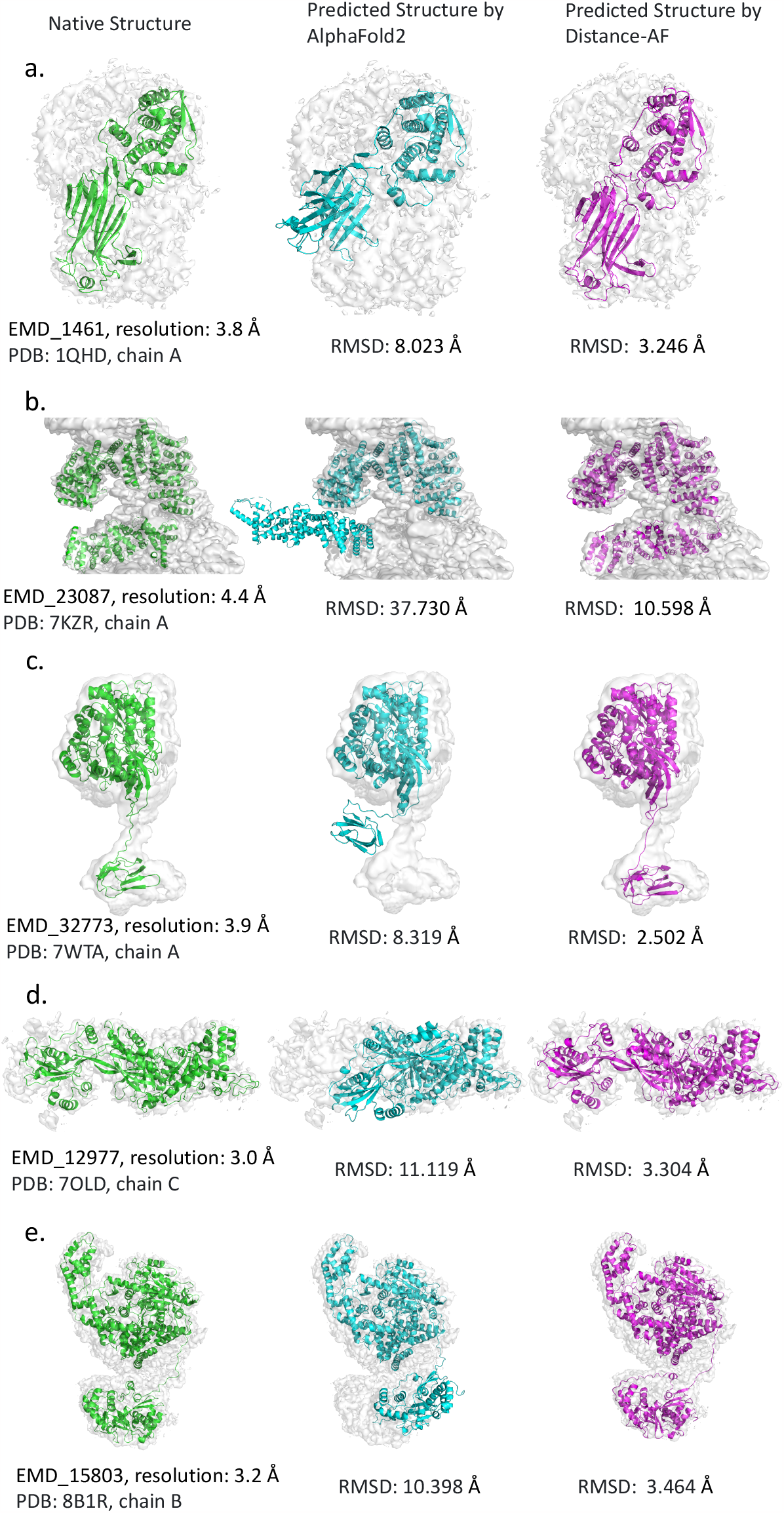
Examples of Cryo-EM deposited structures. The left side shows native structures; the middle part displays initial structures predicted by AlphaFold2, and the right side is final structures predicted by Distance-AF. Specifically, a. Crystal structure of VP6 (PDB:1QHD_A), EMD1461 with resolution of 3.8 Å. The predicted structure by AlphaFold2 results in RMSD of 8.023Å, while Distance-AF improves RMSD to 3.246 Å. b. Cryo-EM structure of EMD 23087, with reported resolution at 4.4 Å(PDB:7KZR_A). Distance-AF improved the structure by achieving much lower RMSD into 10.598 Å by 27.132 Å. c. Cryo-EM structures of human pyruvate carboxylase in apo state with EMD32773 with reported resolution of 3.9 Å (PDB:7WTA_A). Distance-AF successfully lower down RMSD to 2.502 Å. Specifically, in b. and c, those structures are detected as missing residues are masked for better visualization. d. The high-resolution cryo-EM structures of EMD12977 at 3.0 Å (PDB: 7OLD_C). The full atom RMSD on predicted structure by AlphaFold2 is 11.119 Å, in comparison to the one by Distance-AF with 3.304 Å. e. The Structure of EMD15803 at resolution 3.2 Å (PDB: 8B1R_B). Distance-AF predicted at 3.431 Å on RMSD for it but AlphaFold2 produces the structure with RMSD of 10.398 Å.

The first example, illustrated in Figure 5, is derived from PDB 1QHD, specifically from chain A, representing the crystal protein structure of VP6 at a resolution of 3.8 Å. Distance-AF demonstrates notable improvement in the predicted structure, yielding an RMSD of only 3.246 Å. The second example is a target with 1600 residues, identified by the PDB ID 7KZR on chain A, obtained from EMD-23087, with a reported resolution of 4.4 Å. AlphaFold2 exhibits much worse performance in terms of RMSD, resulting in a value of 37.730 Å. A large domain is notably misaligned, nearly extending beyond the designated region. Distance-AF, on the other hand, successfully predicts the large domain within the correct region, thereby rectifying its alignment within the density. This leads to an RMSD value of 10.598 Å. However, it’s worth noting that the relatively elevated RMSD value for the Distance-AF predicted structure in this instance primarily stems from the inaccuracies in local structures predicted by AlphaFold2, an issue that Distance-AF is unable to rectify. The third example, involving chain A from 7WTA, has two distinct domains, corresponding to EMD-32773, with a resolution of 3.9 Å. The structure of human pyruvate carboxylase in the apo state is represented. AlphaFold2’s predicted structures yield an RMSD of 8.319 Å, with the smaller domain falling outside the designated volume, indicating inconsistency with the native structure. In contrast, Distance-AF produces more promising structure, attaining a significantly lower RMSD of 2.502 Å. The fourth example is derived from chain C from PDB 7OLD, deposited from EMD-12977, with a resolution of 3 Å. While AlphaFold2 excels at the individual domain level, one of the domains folds in a manner that compresses towards another, resulting in certain regions mislocating outside the density. This leads to a high RMSD value of 11.119 Å. Distance-AF proves effective in this case by extending the compressed domain to a reasonable level within the volume, thereby lowering the RMSD to 3.304 Å. The final example, selected from chain B of 8B1R, is the DNA binding protein structure of RecBCD in complex with the phage protein gp5.9. This protein is deposited from EMD-15803 with a resolution of 3.2 Å. AlphaFold2 encounters difficulty in predicting one of the domains, which shifts beyond the range of density, resulting in an RMSD of 10.398 Å. In comparison, the structure predicted by Distance-AF demonstrates an RMSD of only 3.431 Å.

### Exploration of Distance-AF to GPCRs

G protein-coupled receptors (GPCRs) are known to undergo structural shifts upon activation or ligand binding, leading to changes in how specific regions within the receptor are arranged in space. These shifts often involve the movement or reorganization of distinct domains or subunits within the receptor, which subsequently bring about functional alterations in signal transduction. In order to thoroughly investigate the potential of Distance-AF, we conducted extensive experiments on GPCRs, with a particular emphasis on the structural changes associated with the transition between active and inactive states, as well as the adjustment of structures to conform to either active or inactive states. Our findings clearly demonstrated that Distance-AF effectively captured these structural alterations and exhibited proficiency in predicting the alternate states. This underscores its effectiveness in leveraging distance constraints to accurately model the dynamic behavior of GPCRs. We established two subtasks to assess the effectiveness of Distance-AF on GPCRs. The first task involved examining state transitions. Given Distance-AF’s proficiency in managing movement, we employed it to facilitate state switching in GPCR targets. This included transitioning between active and inactive states, as well as from inactive to active states. The second task focused on structure refinement. In our examination of individual GPCR targets, we observed instances where AlphaFold2 inaccurately predicted conformational changes. Here, Distance-AF played a crucial role in enhancing the accuracy of predicted structures, specifically in rectifying these mispredictions of conformational shifts.

Subtask 1: Structural rearrangement between native states

We initiated our experiments by focusing on subtask 1, which aimed to facilitate a switch in conformation between active and inactive states. To effectively demonstrate this state transition, we selected two pairs of targets that belong to the same family and share identical sequences. The results are presented in Fig 6.A and 6.B. In Figure 6.A, both targets share the UniProt ID P02699 and are classified under the Rhodopsin family. Chain R of 6OYA represents the active state, while chain A of 3C9L represents the inactive state. Fig 6.A visually indicates the transition from the active state (6OYAR) to the inactive state (3C9LA) facilitated by Distance-AF. Notably, the structural rearrangement is highlighted within the rectangular box. Distance-AF successfully accomplishes this state transition by substituting the cyan chain with the magenta chain, in the magenta chain exhibits a high degree of superimposition with the green, inactive reference structure. Similarly, we conducted tests for the reverse transformation, transitioning from inactive to active states, using a different pair of targets. The active state is represented by chain A of 7BTS, while its inactive counterpart is chain A of 7BVQ. Both targets belong to the family of structures of the Beta-1 adrenergic receptor, sharing the same UniProt ID P08588. Fig 6.B illustrates the results, wherein the magenta structure predicted by Distance-AF shifts from the cyan structure of 7BVQA, which represents the inactive state, towards the green structure, the native active state of 7BTSA. Detailed examination reveals that Distance-AF is able to push TM5 and TM6 outwards, moving them closer to the green structure, thereby aligning with the movement direction observed in the native structure.

**Fig 6.**
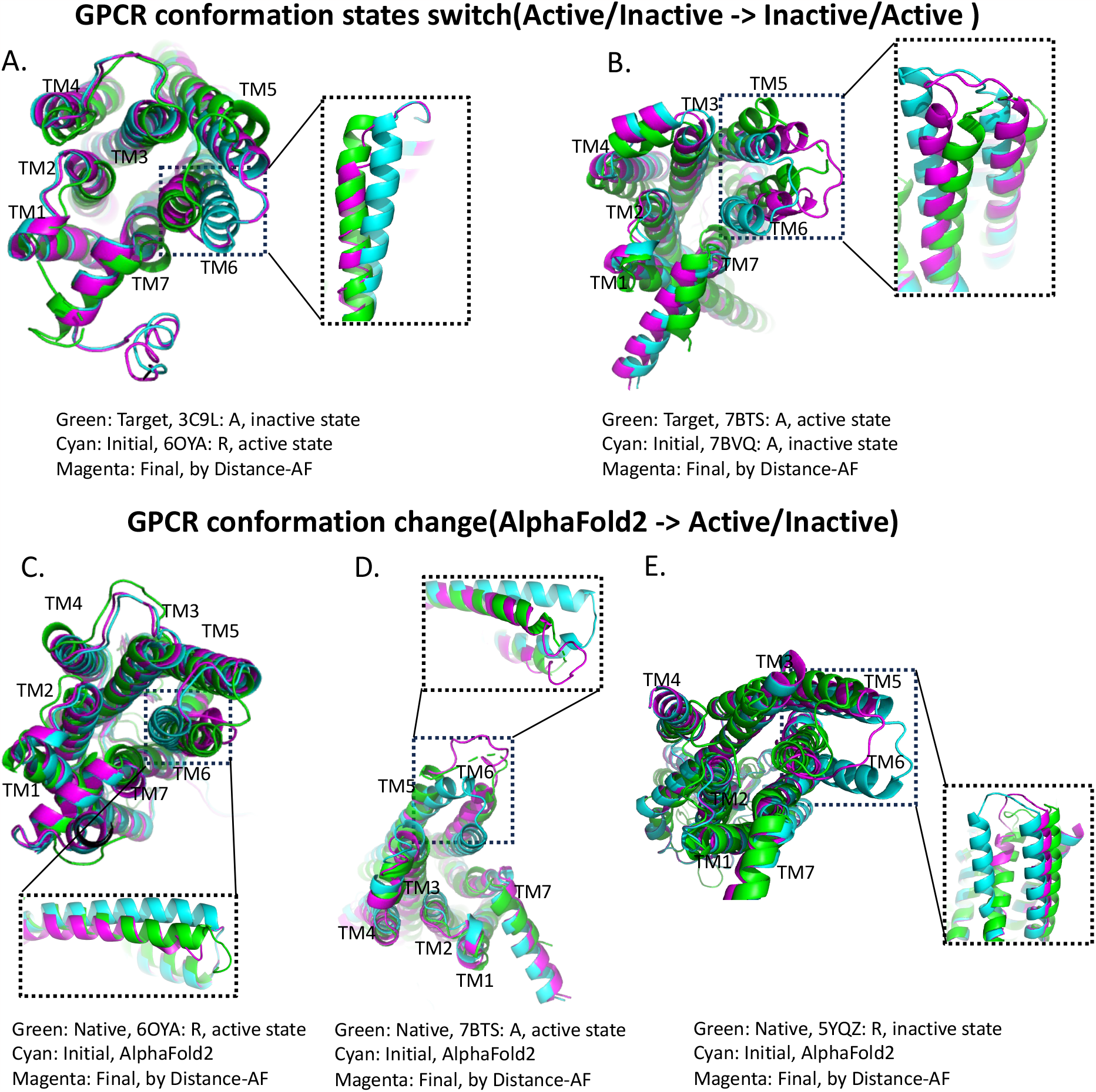
The results of Distance-AF for structural rearrangement on GPCRs. The native structures were depicted in green, representing the reference structures. The cyan structures corresponded to the structures predicted by AlphaFold2, the magenta-colored structures represented the structures predicted by Distance-AF. Upon comparison, it was observed that the structures predicted by Distance-AF exhibited notable improvements and refinement compared to the initial predictions by AlphaFold2 on specific conformation changes of GPCRs. The magenta structures displayed enhanced agreement with the native structures, indicating the success of Distance-AF in effectively capturing and rectifying structural discrepancies.

Subtask 2: Correction of structural rearrangement on AlphaFold2 structures

Next, we focused on subtask 2 by selecting three targets with the objective of rectifying erroneous conformations in the predictions generated by AlphaFold2. Among these, two of the targets, namely 6OYAR and 7BTSA, are in an active state, while the third, 5YQZR, is in an inactive state. This allows for a comprehensive assessment of Distance-AF’s performance in both active and inactive state corrections, as depicted in Figures 6.c, 6.d, and 6.e. In the case of active states for both 6OYAR and 5YQZR, Distance-AF exhibits a notable outperformance over AlphaFold2. It accurately predicts the conformational changes, aligning closely with the native structure. These details are clearly illustrated in Figures 6.c, 6.d, and 6.e for both states. More details can be observed in the rectangle areas, where magenta (the structures by Distance-AF) shows more agreement to green (native structures) than cyan (AlphaFold2 predicted structures).

### Application of Distance-AF to NMR structures

NMR (Nuclear Magnetic Resonance) is a powerful method to investigate the dynamics of biological molecules in solution. Those solution-state structures, exploring the conformation of the molecule in a liquid (typically water) environment. Therefore, NMR structures are often represented as an ensemble of diverse conformers. However, existing methods even accurately like AlphaFold2 are limited to predict close conformers per target, while other diverse conformers are co-existed not getting predicted. Distance-AF is able to address the problem by applying a groups of distance constraints shared by different native conformers to predicting corresponding conformers in consistent with the given distance constraints. In this case, Distance-AF achieves the objective that distinguishable conformers of the same target can be predicted accurately. To evaluate how Distance-AF performs on NMR structures with multiple conformers, we carefully picked out 3 targets with more than 20 conformers to test, 2M8P, 1DMO and 1TNW. The PDB entry 2M8P[26], presents the solution structure of the HIV-1 capsid protein, which exists as a dynamic equilibrium between monomeric and dimeric forms among 100 conformers. Besides, 1DMO and 1TNW are solution structure of calcium binding proteins, where 1DMO is *Ca*^*2+*^ free calmodulin with 30 conformers compared to 1TNW is *Ca*^*2+*^ saturated with 23 conformers. Even though 2 targets share similar intra-domain local structures, the presence of leads to major variations to domain surfaces, and domain conformation changes.

Given that common distances between amino acid pairs are typically limited, usually fewer than three, while encompassing over fifteen conformers per target, the Distance-AF employs approximately common distance constraints, set at a cutoff of 2 Å. These constraints are derived from manually selected conformers, each structurally distinct from the others, yet sharing several pairs of residues with distances within a difference of 2 Å. We illustrate the examples of Distance-AF’s performance on NMR structures in Fig. 7. For the left side of all panels, the selected native conformers are shown, in comparison to the predicted conformers by Distance-AF are at right side. The first target (Fig. 7a) exemplifies a calcium-saturated structure in solution, with eight conformers selected from a pool of twenty-three to establish a set of eight distance constraints. Distance-AF successfully generates seven reasonable structures out of eight, demonstrating an average RMSD of 4.629 Å *a*gainst their corresponding native structures. The second one (Fig. 7b), comprises thirty conformers with calcium absence in the solution. Among the seven selected native conformers, five are accurately predicted by Distance-AF, yielding an average RMSD of 4.981 Å. The last target (Fig. 7c) represents the solution structure of the HIV-1 capsid protein, encompassing one hundred conformers. Distance constraints applied to Distance-AF originate from twelve conformers, with seven out of twelve being accurately predicted, resulting in an average RMSD of 4.871 Å.

**Fig 7.**
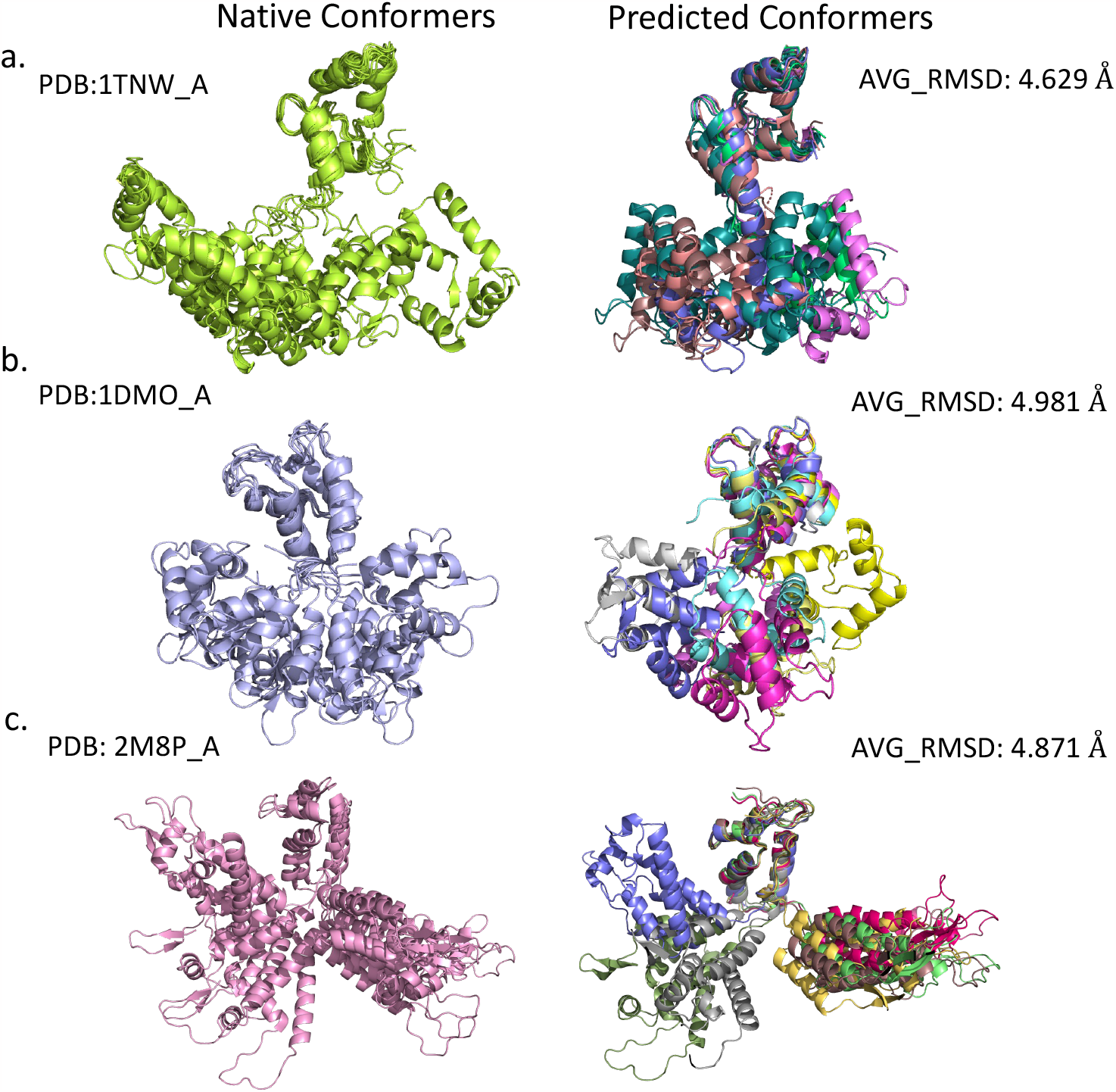
Examples of multiple conformers. The selected native conformers are shown at left, which shares the same group of distance constraints, while the conformers predicted by Distance-AF are shown at right side. a. chain A of 1TNW, NMR solution structure of calcium injected skeletal muscle troponin C reported with 23 conformers, Distance-AF achieves average full-atom RMSD across 7 predicted conformers is 4.629 Å; b. chain A of 1DMO, the solution structure of apo calmodulin with the removal of *Ca*^*2+*^ with 30 conformers reported. Across the 5 predicted conformers, the average full-atom RMSD is 4.981 Å; c. chain A of 2M8P, the structure of the W184AM185A mutant of the HIV-1 capsid protein getting 100 conformers, Distance-AF predicted 7 conformers with average RMSD at 4.871 Å.

**Fig 8.**
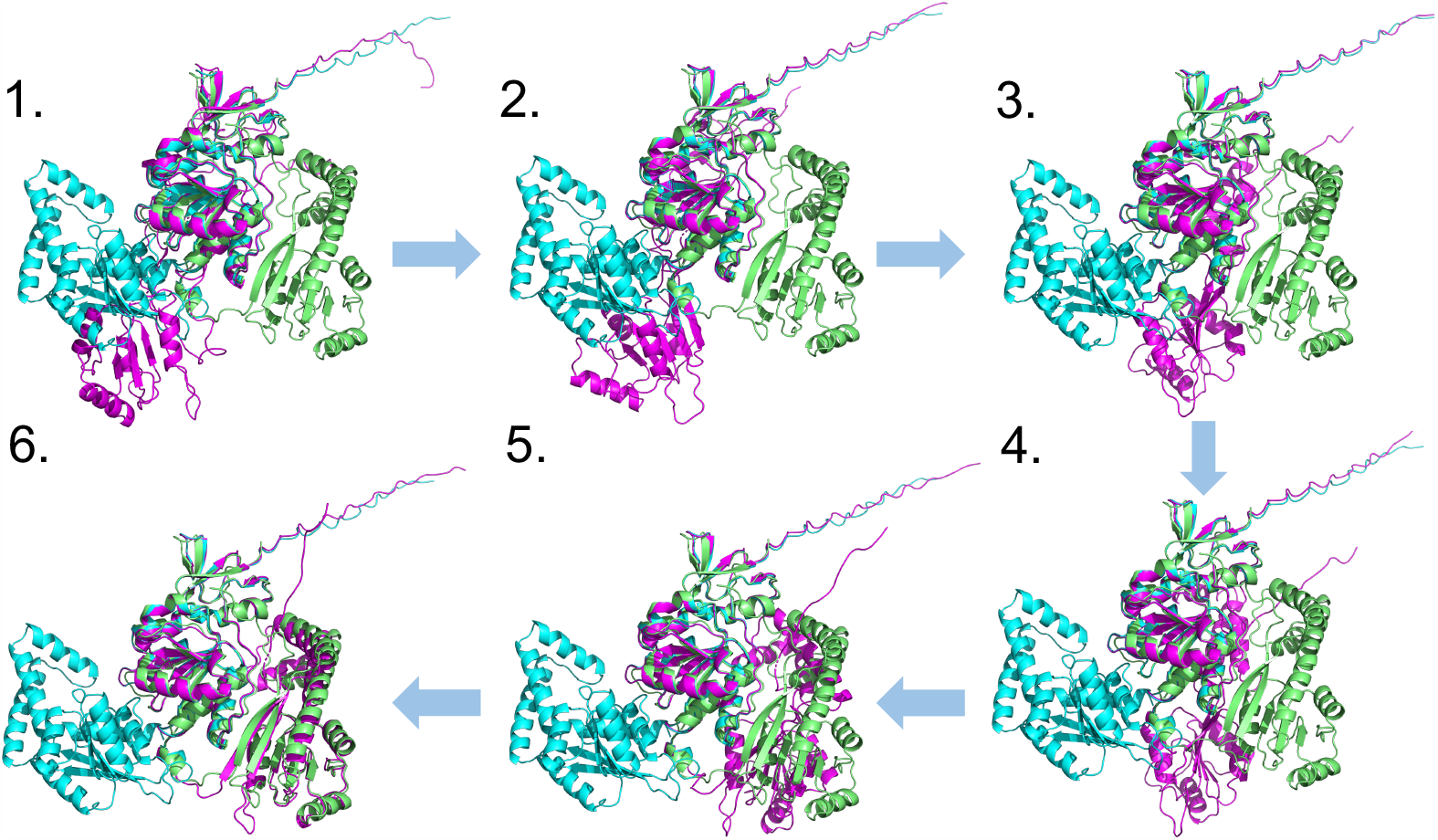
The ability of Distance-AF by capturing domain motion during different stages is illustrated in the predicted structures of PDB 6P66 above. The native structure is shown in green, while the starting structure predicted by AlphaFold2 is shown in cyan, the 2 structures are always fixed. The predicted structure by Distance-AF is shown in magenta, where one of its domains moves gradually from left top (stage 1) to left bottom (stage 6).

## Discussion

We propose Distance-AF, a novel deep learning method that harnesses the power of the AlphaFold2 framework while incorporating distance constraints to improve protein structure prediction. These constraints serve as essential spatial information regarding the proximity and relationships between specific residues and atoms within the protein structure, which can be derived from experimental techniques such as cryo-electron microscopy (cryo-EM), nuclear magnetic resonance (NMR) spectroscopy, or chemical crosslinking. In Distance-AF, the given distance constraints are integrated into the training process, navigating protein folding by simultaneously optimizing its predictions to satisfy both the fundamental principles of protein structure and the experimentally derived distance constraints. Through the benchmarking experiments, Distance-AF show the potential to generate more accurate models compared to AlphaFold2 and Rosetta. Meanwhile, they exhibited exceptional performance on challenging protein classes such as G-protein-coupled receptors (GPCRs), structures obtained through Cryo-EM and multiple conformation achievements on NMR targets. Those advancements presented by Distance-AF have significant implications for various fields, including drug discovery, protein engineering, and understanding the molecular mechanisms underlying biological processes. However, it’s important to note a limitation that Distance-AF’s applicability is currently optimized for only protein monomers with distinct domains. For much more flexible and uncompact structures, Distance-AF is unable to achieve good performance. It is conceivable that future iterations of Distance-AF may extend its applicability to protein multimers or even much larger protein complexes, further broadening its scope and impact across wider biological system applications.

## Supplementary Information

### Example of Domain Motion Trajectory

To provide a more comprehensive understanding of how Distance-AF effectively corrects the incorrect domain shifts observed in AlphaFold2’s predictions, Fig 7 shows the dynamic movement of domains at various stages during the training process. This figure offers a visual representation of the domain correction mechanism employed by Distance-AF.

### Structural changes with learning curves

To obtain insight into the evolution of the predicted structure over 1000 epochs during the overfitting process, Fig 9 illustrates the structural changes at distinct intervals. Specifically, we selected significant time points representing various stages across four distinct loss functions. Epoch 0 denotes the initial structure, while epoch 7 marks the peak of the violation loss. At epoch 100, the distance loss is in a transitional phase of reduction, while epoch 211 represents the point where the distance loss reaches its minimum. Epoch 255 corresponds to a stage where both Fape and angle loss attain their highest values. At epochs 400 and 600, Distance-AF is engaged in the recovery of local structures, indicated by the declining Fape and angle loss values. Finally, epoch 1000 shows the ultimate predicted structure.

**Fig 9.**
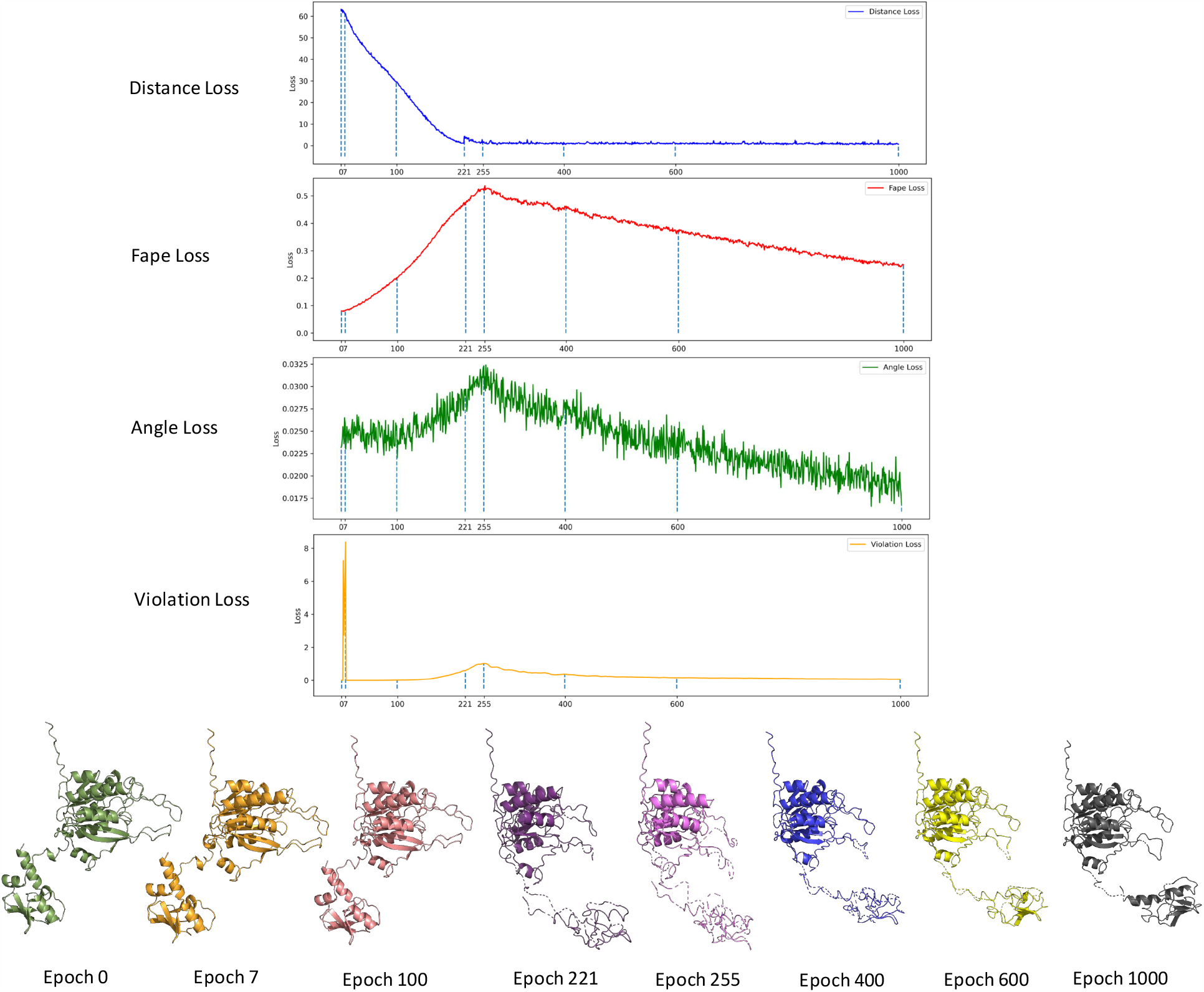
The structural changes within 1000 epochs with evolutional learning curves.

### Correlation study about distance loss to RMSD and TM-Score

To further assess whether the proposed distance loss function aids in directing the domain towards the desired domain orientation, in accordance with the goals of RMSD and TM-Score to achieve more accurate structure, Fig 10 illustrates the correlation between the distance loss value at the last epoch and the final full-atom RMSD value in a., as well as the final TM-Score in b. Fig 10.a demonstrates the relationship between the absolute value of distance loss and full-atom RMSD across all 25 targets, while Fig 10.b illustrates the correlation between distance loss and TM-Score. When the distance loss value is relatively low, it signifies that the predicted structure adheres more closely to the provided distance constraints. Consequently, the corresponding RMSD value is also relatively lower, and the TM-Score is higher, indicating the folded structure agrees more with native structure. Both correlations indicate that the defined distance loss works consistently with the optimization of structural folding.

**Fig 10a.**
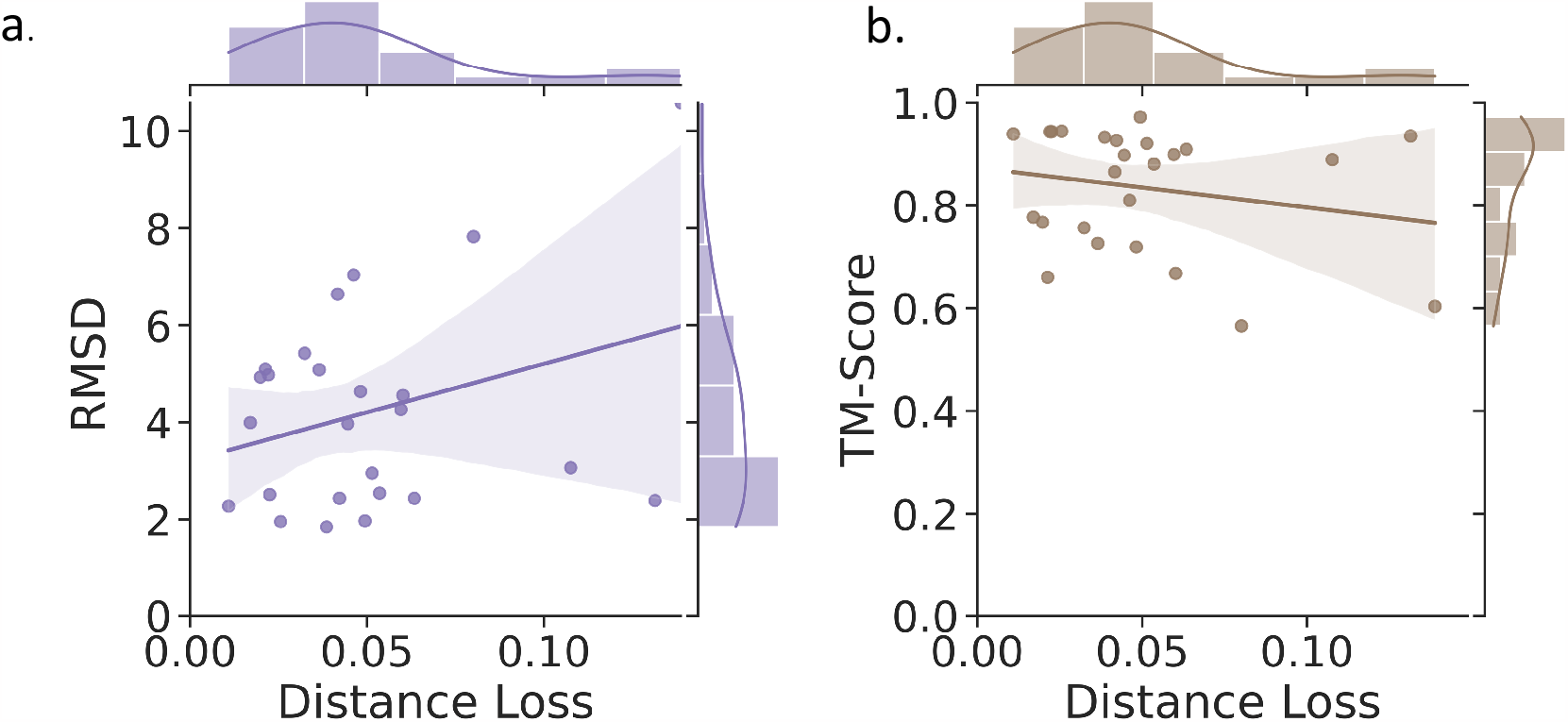
The correlation on distance loss(x-axis) to full-atom RMSD(y-axis); b. The correlation on distance loss(x-axis) to TM-Score(y-axis)

## Availability

The code of Distance-AF program is available at https://github.com/kiharalab/Distance-AF.

## Acknowledgments

This work was partly supported by the National Institutes of Health (R01GM133840 and 3R01GM133840-02S1) and the National Science Foundation (CMMI1825941, MCB1925643, IIS2211598, DMS2151678, DBI2146026, and DBI2003635).

